# Easily phylotyping *E. coli* via the EzClermont web app and command-line tool

**DOI:** 10.1101/317610

**Authors:** Nicholas R. Waters, Florence Abram, Fiona Brennan, Ashleigh Holmes, Leighton Pritchard

## Abstract

The Clermont PCR method of phylotyping *Escherichia coli* has remained a useful classification scheme despite the proliferation of higher-resolution sequence typing schemes. We have implemented an in silico Clermont PCR method as both a web app and as a command-line tool to allow researchers to easily apply this phylotyping scheme to genome assemblies easily.

**Availability and Implementation:** EzClermont is available as a web app at http://www.ezclermont.org. For local use, EzClermont can be installed with pip or installed from the source code at https://github.com/nickp60/ezclermont. All analysis was done with version 0.4.0.

**Supplementary information:** Table S1: test dataset; S2: validation dataset; S3: results.

---

*Escherichia coli* is among the most widely studied organisms, and the species is very diverse [11, 13]. Because of this diversity, many methods have been developed to differentiate the different *E. coli* lineages. In 1987, Selandar and colleagues used electrophoretic analysis of a 35 enzyme digest to classify the *E. coli* Reference Collection (ECOR) in 6 phylogenetic groups (A-F) [13]. Clermont and colleagues published their triplex PCR method [4] of phylotyping, which proved to be an extremely valuable tool to differentiate groups A, B1, B2, and D, being cited over 625 times as of April 2018. In 2013, Clermont and colleagues published an update to this work [5], in which they showed that by adding a 4th set of primers (with additional primers to differentiate the subgroups and the cryptic clades), higher resolution could be achieved, as this expanded the method to detect groups E, F, and differentiate the cryptic clades. This approach has been widely adopted, as the method is reliable, easy to interpret, and can be performed rapidly.

Other sequence typing schemes have been developed to classify *E. coli* strains. These include the Achtman 2012 7 gene Multi Locus Sequence Typing (MLST) [1, 2], Michigan EcMLST [12], whole-genome MLST (http://www.applied-maths.com/applications/wgmlst), core-genome MLST [7], two-locus MLST [17], and ribosomal MLST [10]. All these methods classify *E. coli* with greater accuracy and granularity than the phylotyping, but at the cost of interpretability. The Clermont 2013 phylotyping scheme remains a regularly utilised tool in classifying *E. coli*.

We developed EzClermont to provide a simple implementation of the Clermont phylotyping algorithm to genome assemblies. For researchers unfamiliar with command-line tools, we have implemented the software as a web application; for those needing to process large numbers of assemblies, a command-line interface can be installed via pip.

In short, the software uses constrained string matching as an in silico PCR to determine the presence or absence of the alleles used to determine the phylotype. As assemblies may contain alleles interrupted by breaks between contigs, we give the user the option to allow partial matches (ie, if one of the two primers matched, but the expected position of the other primer fell beyond the sequence end).

As PCR primers do not necessarily need 100% sequence identity to function, we determined the variability at the priming sites in 523 strains. To do this, we downloaded the genome assemblies from NCBI Bioprojects PRJNA218110, PRJNA231221, and PRJNA352562. From each assembly, we extracted the 7 regions matching the theoretical amplicons of the quadriplex, E-specific, C-specific, and E/C control primer sets from Clermont 2013. Any differences between a sequence and the primer sequence reported in Clermont 2013 were incorporated into the search query, except for differences in the last 5 nucleotides on the 3’ regions (as those can be used to differentiate alleles) [15].

To assess the performance of EzClermont, we selected a test dataset and a validation dataset. Additionally, the strains from Clermont, 2013 Figure 1 are used as unit tests in the package.

**Figure 1:**
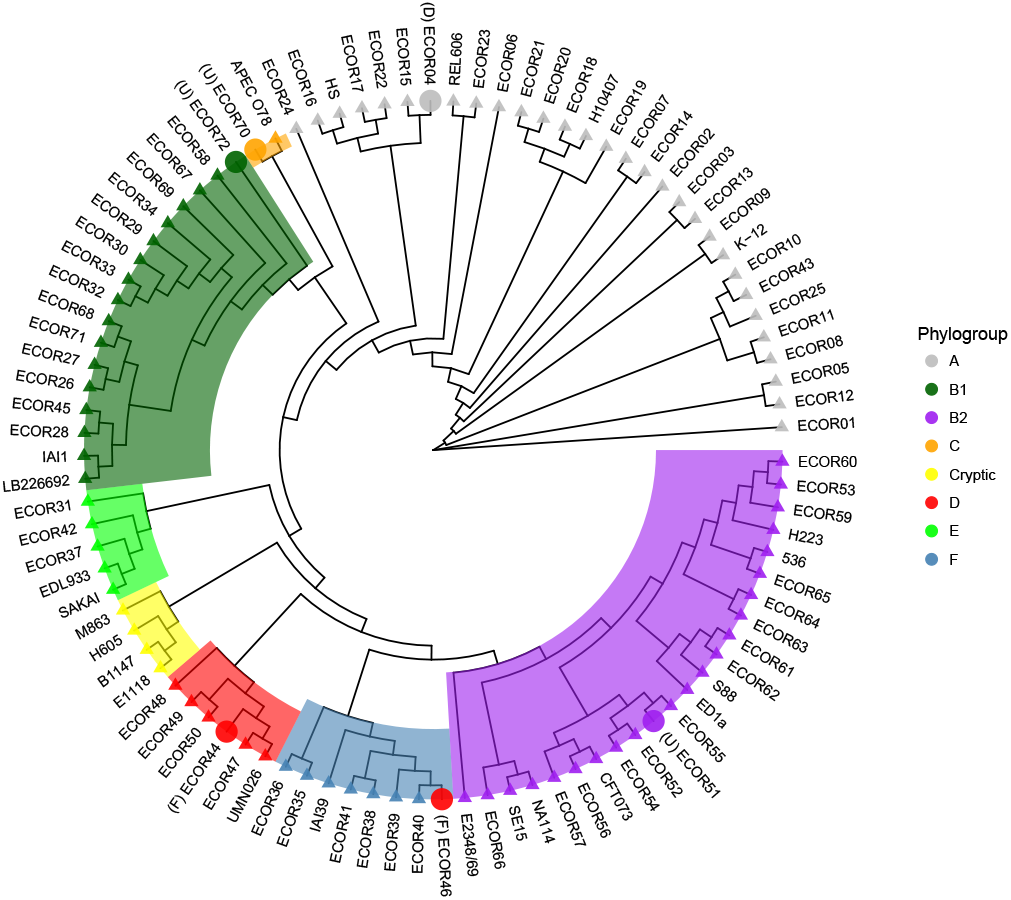
Parsimony cladogram of strains from Clermont et al 2015. Tree was generated with kSNP3 (k=19). Enlarged circular tips show where EzClermont differed from reported phylogroup (EzClermont type show in brackets).

As a test set, we used strains listed in Sims and Kim 2011 [14] (Table S1), and the validation set of 95 strains was the genomes from Clermont 2015 [6] (Table S2)^1^. Comparing the reported phylogroup and the EzClermont phylogroup for the 19 strains in Sims and Kim (excluding strains reported in both Clermont 2015 and Sims and Kim), 3 of the 19 did not agree, but two of those (IAI39, SMS-3-5) were shown by other works to have the phylotype that EzClermont predicted (see Table 1). The one strain that typed differently (APEC01) was examined and was found to have the ArpA allele that is not normally detected in B2 strains.

**Table 1:**
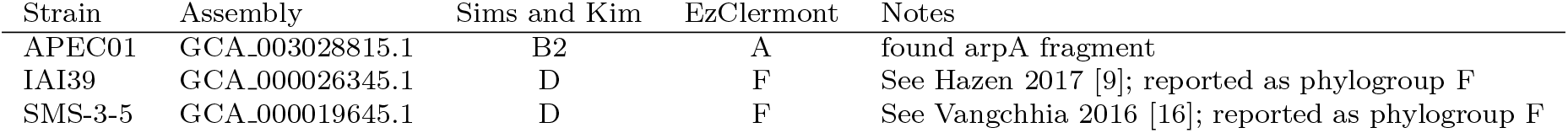
Comparing EzClermont to phylotypes reported by Sims and Kim 2011 [14]

We ran EzClermont on the 95 strains from Clermont 2015 and compared the results to the reported phylotype; 89 of the 95 strains classifications matched. To determine whether the inconsistent phylogroup assignments matched phylogeny, we then generated a parsimony tree using kSNP3 [8], and plotted with ggtree [18]. This revealed that the EzClermont classification of ECOR46 (similar IAI39 and SMS-3-5) appears to match the true phylogeny, as opposed to the phylogroup reported in the literature (Figure 1). Of those that didn’t match, all detected at least one theoretical amplicon that was not reported to be there (Table S3). Further, a wide application of EzClermont by Zhou et al. [19] to representative *E. coli* strains in Enterobase was largely in agreement with both higher-resolution sequence typing and with ClermonTyping [3], another in silico tool for detecting Clermont types.

Considering both the testing and validation datasets (114 strains), EzClermont has an accuracy of 94%. Given the ease of use of the web app for simple queries, its incorporation into Enterobase, and the standalone speed of execution for larger batches, we hope that EzClermont will be of continued use to the community.

## Acknowledgements

Many thanks to Stephen Nolan, Dr. Corine Nzeteu, and Dr. Alma Siggins for their comments on the manuscript.

## Footnotes

1 6 of the 101 total strains were omitted as no genome assembly was available.

